# Unilateral striatal deep brain stimulation improves cognitive control

**DOI:** 10.1101/2025.11.10.686309

**Authors:** Elizabeth M Sachse, Evan M Dastin-van Rijn, Jonathan P Bennek, Michelle C Buccini, Megan E Mensinger, Francesca A Iacobucci, Adriano E Reimer, Alik S Widge

**Affiliations:** Department of Psychiatry & Behavioral Sciences, University of Minnesota, 606 24th Ave S, Minneapolis, MN 55454; Department of Neuroscience, University of Minnesota, Jackson Hall, Room 6-145, 321 Church St SE, Minneapolis, MN 55455; Department of Biomedical Engineering, University of Minnesota, Nils Hasselmo Hall, 7-105, 312 Church St SE, Minneapolis, MN 55455

**Author notes:** **Corresponding Authors:** Elizabeth M Sachse, Alik S Widge.

## Abstract

Deep brain stimulation (DBS) of the ventral capsule/ventral striatum (VCVS) can treat obsessive-compulsive disorder (OCD) and other psychiatric conditions. Yet, optimizing its clinical efficacy is a major challenge, often hindered by incomplete knowledge of how stimulation parameters and targets affect neural activity and behavior. VCVS DBS is thought to work in part by improving cognitive control, an important decision-making component that is impaired in OCD and other illnesses. The magnitude of this cognitive control enhancement was shown to be lateralized, with right-unilateral stimulation being the most effective. Prior work developed a preclinical model of VCVS DBS by leveraging the cognitive control construct, which can be measured in humans and rodents and is modulated by analogous brain circuits. However, this work did not address laterality effects observed in humans or examine left/right stimulation differences. These effects may be critical for maximizing therapeutic benefit while avoiding aversive outcomes. This study aimed to investigate lateralization in the rodent model, where bilateral stimulation of the mid-striatum was previously shown to improve cognitive control. Right and left-unilateral stimulation reduced response times without changing accuracy, replicating the cognitive control improvement from bilateral stimulation. With computational modeling, we show that bilateral and unilateral stimulation modifies the same decision-making variables to drive this behavior change. We also establish that females have the same cognitive control improvement from stimulation as males. These findings increase our understanding of cognitive control circuits and strengthen the validity of the rodent model as a translational platform to study VCVS DBS’s therapeutic mechanisms.

**Significance Statement:** Here, we demonstrate that stimulating just one side of the brain (e.g. unilaterally) can be as effective as bilateral stimulation for improving cognitive control, the ability to adjust thoughts and decisions in response to environmental changes. These findings in rodents match results from prior human deep brain stimulation (DBS) studies, highlighting the validity of this preclinical model to study DBS’s therapeutic mechanisms. We hypothesize that unilateral stimulation may be preferable to maximize cognitive benefits without causing off-target effects, while also reducing surgical invasiveness. Further, we demonstrate that females have the same cognitive control improvement from stimulation as males. Overall, this work answers important outstanding clinical questions regarding laterality and sex in DBS therapies for psychiatric illnesses.

## Introduction

Deep brain stimulation (DBS) delivered to the ventral internal capsule/ventral striatum (VCVS) is approved for treatment-refractory obsessive-compulsive disorder (OCD) (Widge et al., 2019b; Denys et al., 2020; Tyagi et al., 2022), and is under investigation for major depressive disorder (MDD) (Widge et al., 2019b; Runia et al., 2023), and other conditions (Lipsman et al., 2013; Bach et al., 2023). However, DBS’s clinical potential is limited by an incomplete understanding of how it modifies neural networks (Sullivan et al., 2021; Widge, 2024). This makes stimulation targeting and optimization challenging, especially given patient heterogeneity (Karas et al., 2019; Hitti et al., 2023). Traditional readouts of treatment efficacy include successful electrode placement and self-reported symptoms. However, patient outcomes vary widely (Li et al., 2020; Graat et al., 2022), and self-reported symptoms are subjective and unreliable (Sullivan et al., 2021; Widge, 2024). Animal models allow for circuit-specific approaches and controlled environments, but cannot adequately model diagnoses like “depression” (Monteggia et al., 2018; Redish et al., 2021).

A potentially more productive approach to studying DBS mechanisms is to measure underlying cognitive constructs that are impaired in psychiatric conditions (Insel et al., 2010; Robbins et al., 2012, 2024; Cuthbert and Insel, 2013). For example, OCD, MDD, anxiety, and addiction all involve dysfunctional cognitive control, the ability to adjust thoughts and behaviors in response to a changing environment (Waltz, 2017; Kim et al., 2019; Uddin, 2021; Egner, 2023; Grant and Chamberlain, 2023; Widge, 2024). Cognitive control is implemented via cortico-striatal-thalamo-cortical (CSTC) circuits, the same circuitry engaged by VCVS DBS (Smith et al., 2019; Widge et al., 2019a; Menon, 2020; Basu et al., 2023; Badre, 2024; Sachse and Widge, 2025).

With these insights, recent studies have assessed the effects of VCVS neurostimulation on cognitive control. In OCD and MDD patients, as well as patients without diagnosed psychiatric illness, VCVS DBS reduced response times (RTs) without increasing errors on a behavioral task, signaling improved cognitive control (Widge et al., 2019b; Basu et al., 2023). Stimulation also strengthened theta oscillations in dorsolateral prefrontal cortex (PFC), an area engaged during cognitive control, indicating a potential mechanism by which DBS exerts its effects. This findings were replicated in subsequent studies (McLaughlin et al., 2021; Allawala et al., 2023). One group tested whether stimulation of sub-regions within the VCVS differentially affected cognitive control. The magnitude of both the behavioral improvement and PFC theta change varied by electrode contact, with the right dorsal contact being the most effective (Basu et al., 2023). This laterality effect suggests a “sweet spot” for stimulation along the dorsal-ventral and medial-lateral axes of the VCVS. However, these results, like many human DBS studies, are limited by small patient numbers and clinical experimental constraints.

To address these limitations, a reverse translational rodent model of VCVS DBS was created (Reimer et al., 2024). Based on human-rodent CSTC circuit homology (Heilbronner et al., 2016; Coizet et al., 2017), different possible analogs of the human VCVS were stimulated during a cognitive control task (Park et al., 2016; De Oliveira et al., 2021). DBS-like stimulation of the mid-striatum replicated the reduced RTs observed in humans. Further, the rodent behavioral effect followed a comparable anatomical gradient to humans (Basu et al., 2023). Computational modeling showed that stimulation increases cognitive control through similar behavioral mechanisms in both species, validating the animal model (Basu et al., 2023; Reimer et al., 2024).

A missing component in that animal model was an examination of laterality and comparison to differential effects of left/right VCVS stimulation in humans. Laterality effects are common in DBS, and in neuroscience generally (Hershey et al., 2008; Lueken et al., 2008). Variation in the effectiveness of different DBS electrode contacts and anatomical targets has been frequently observed in DBS patients. (Sturm et al., 2003; Conroy et al., 2021; Lin et al., 2021). Therefore, laterality effects may be critical to DBS’s therapeutic benefits, particularly for psychiatric conditions where we may want to boost certain cognitive functions without impairing others. Most psychiatric DBS interventions target both hemispheres, then activate different contacts or combinations of contacts to try to find each patient’s ideal configuration. (Hamani et al., 2014; Widge and Dougherty, 2022). However, for some targets, unilateral DBS may provide equivalent symptom relief as bilateral DBS, (Sturm et al., 2003), with fewer aversive side effects. (Widge et al., 2016; Provenza et al., 2021). These laterality effects warrant further investigation but are difficult to study in humans given limitations mentioned above.

Here, we investigated laterality effects within the mid-striatal DBS model. If this model is truly representative of VCVS DBS, then unilateral stimulation, particularly right-sided, should improve cognitive control similarly to bilateral stimulation. We compared how bilateral, right-unilateral, and left-unilateral stimulation affected RTs versus no stimulation (“OFF”) on a cognitive control task. We found that unilateral and bilateral stimulation similarly reduce RTs, with no significant differences between right and left stimulation. Building on prior work, we also show stimulation enhances cognitive control similarly in females and males.

## Methods

### Animals

17 adult Long-Evans rats (9 male, 250-400g; 8 females, 200-350g) were obtained from Charles River Laboratories (Wilmington, MA) and housed on a 12:12 dark/light cycle. After acclimation, rats were handled for 5 min/day for at least 4 consecutive days to familiarize them with the experimenters. To motivate task performance, rats were food-restricted to maintain 85-90% of their original body weight, with supplemental food provided if task rewards were insufficient to meet their caloric needs. Rats were at least 3 months old prior to any food restriction. Rats were given ad libitum access to standard food for at least 7 days before undergoing stereotaxic surgery. All experiments were approved by the University of Minnesota Institutional Animal Care and Use Committee (protocol 2104-39021A) and complied with NIH guidelines.

### Set-Shifting Task

To measure cognitive control, we used a Set-Shift task (Figure 1A) modified from previous studies, including studies specifically focused on cognitive control and brain stimulation (Darrah et al., 2008; Park et al., 2016; De Oliveira et al., 2021; Reimer et al., 2024). The task occurred in operant chambers (25 x 29 x 25cm; Coulbourn Instruments, Holliston, MA), containing three illuminable nose-poke holes with infrared detectors, a food dispenser on the opposite wall (45mg sucrose, BioServ, Flemington, NJ), and an overhead video camera, enclosed in sound-attenuating boxes (Med Associates, Chicago, IL). Task stimuli were controlled by GraphicState 4.0 (Coulbourn Instruments) or *pybehave/OSCAR* software (Dastin-van Rijn et al., 2023).

**Figure 1:**
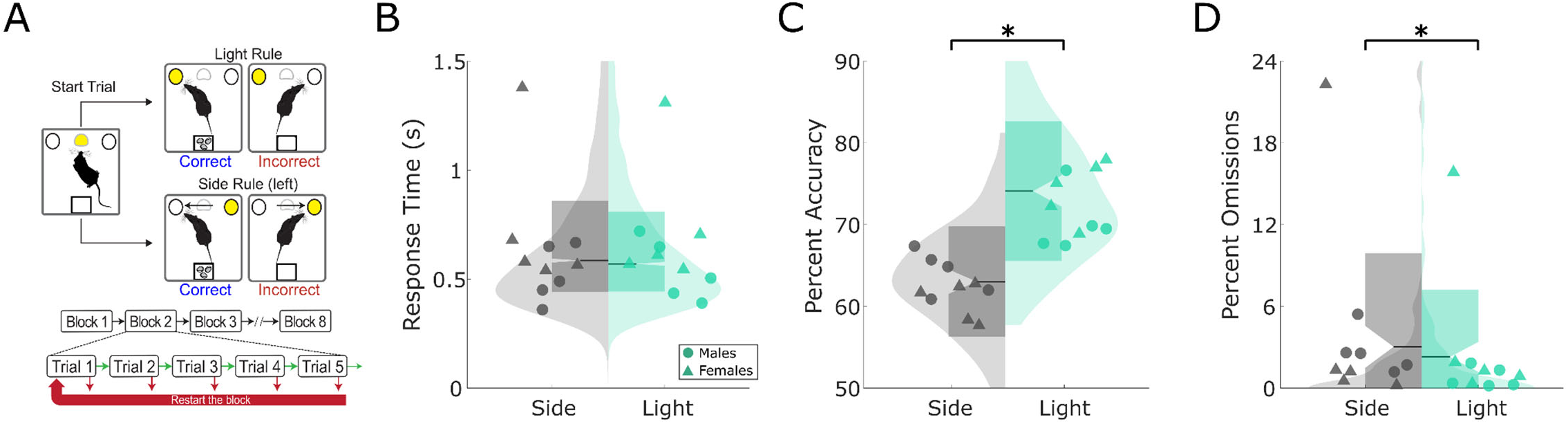
Using a Rodent Set-Shift Task as a Translational Model of Cognitive Control. A) Set-Shift task. Rats initiate a trial by poking the middle nosepoke and then must nosepoke either an illuminated hole (Light rule) or ignore the light and always nosepoke on a specific side of the chamber (Side rule). The rules are not cued, and shift to a new rule after the rat has sequentially completed 5 correct trials of the current rule. Errors reset to the beginning of the current 5-trial block. B) Raw RT for non-stimulated sessions by rule type. Scattered points are individual rats’ median RTs. RTs were not significantly different for side vs. light rule trials (p = 0.0774). C) Raw accuracy percentage for non-stimulated sessions, by rule type. Scattered points are individual rats’ mean accuracy percentages. Rats were more accurate on light rule trials (right) compared to side rule trials (left) (*, p<0.00001). D) Raw percent omissions for non-stimulated sessions by rule type. Scattered points are individual rats’ mean percent omissions. Rats omit more side rule trials than light rule (*, p = 0.000017).

Rats learned and adjusted their responses between a cue-driven “Light” rule and a spatial “Side” rule. Rats had to poke the illuminated middle nose-poke to initiate a trial, then poke one of the two peripheral nose-pokes, one of which was illuminated. Rats were reinforced with a single reward pellet for each correct response. The Light rule required rats to poke the illuminated nose-poke hole, regardless of its spatial location. The Side rule required rats to poke a specific spatial location (left or right), regardless of which one was illuminated. The Light rule is typically easier to execute because it requires a cue driven response to a visually salient stimulus (Figure 1B and C). After rats make five consecutive correct choices, the rule switches to the other dimension, requiring rats to shift their behavior to continue receiving rewards. No cue was provided, besides the absence of a reward, to signal rule changes. A block was defined as the sequential trials on a single rule, regardless of accuracy. The task contained eight blocks, requiring the rats to shift seven times per test session. After reaching testing criteria (see Figure 2B and (De Oliveira et al., 2021)), rats were implanted with stimulating electrodes.

**Figure 2:**
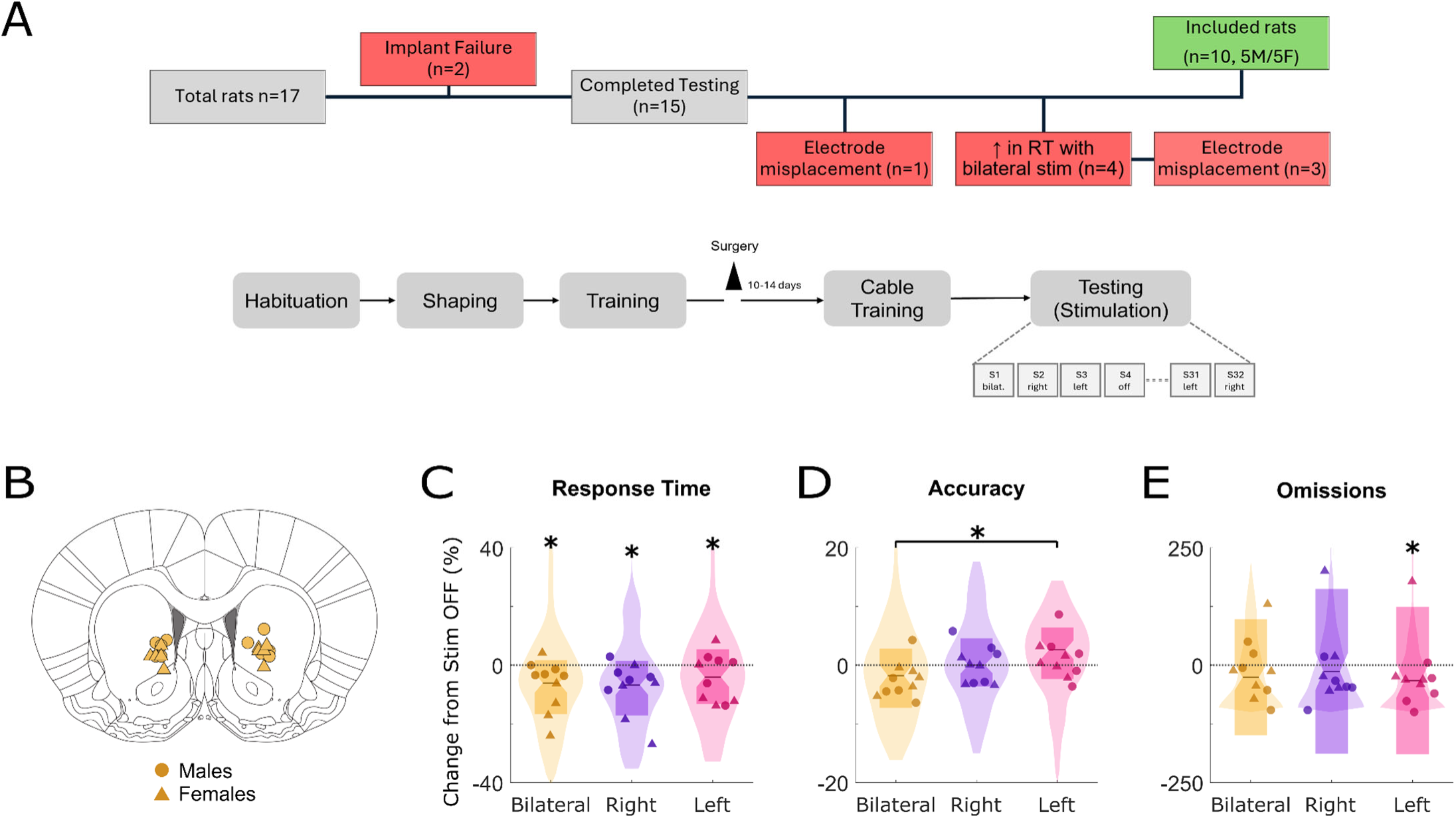
Unilateral mid-striatal stimulation replicates bilateral stimulation’s enhancement of cognitive control. A) CONSORT diagram and experimental schedule diagram. Rats with significant increases in RT with bilateral stimulation and/or electrode mistargeting were excluded from the study. B) Location of electrodes for electrical stimulation of the mid-striatum. Medial-lateral and dorsal-ventral coordinates were plotted onto the rat atlas coronal section that corresponds to the average anterior-posterior coordinate. C) Change in RT with bilateral (gold) or unilateral (right: purple, left: pink) stimulation as percent-change from OFF RT. Distributions quantify median percent-change in RTs across Set-Shift sessions, and individual scatter points represent median percent-change in RT of each rat with bilateral or unilateral stimulation. **\*** indicates significant stimulation-condition intercept from gamma-distributed GLM. D) Change in accuracy with bilateral or unilateral stimulation as percent-change from OFF accuracy. Distributions quantify mean accuracy rates across Set-Shift sessions, and individual scatter points represent mean accuracy percentages of each rat. E) Change in trial omissions with bilateral or unilateral stimulation as percent-change from OFF omissions. Distributions quantify mean omission rates across Set-Shift sessions, and individual scatter points represent mean omission rates of each rat.

After recovery from surgery, rats underwent 32 testing sessions (see “Electrical Stimulation section below). Test sessions consisted of four protocols with different rule orders to prevent predictability of the next rule/trial. Each protocol started and ended with a sequence of 10 trials, in which rewards were given randomly, independent of the rats’ choice to ensure the rat had no a priori rule information before the start of the first rule block. Rule sequences within protocols were counterbalanced so that all possible rules were presented with equal frequency. Protocol order was balanced with stimulation conditions, and was the same for each rat, except when a session could not be run (an equipment malfunction or abnormal testing conditions). Protocols missed due to these conditions were appended to the end of the remaining sessions.

### Surgery for bilateral stimulating electrodes

Rats were anesthetized with isoflurane, given preoperative analgesia, and mounted in a stereotaxic frame. The tissue was resected over target areas and craniotomies were drilled for anchor screws and electrodes. Next, bipolar twisted platinum iridium insulated electrodes (MS333/8-BIU/SPC, Plastics One, Roanoke, VA), coated with Vybrant DiI or DiD cell labeling solution (Invitrogen, Eugene, OR) were placed bilaterally in the mid-striatum (+1.4 AP, ±2.0 ML, -5.8 DV, target area based on (Reimer et al., 2024)). Electrodes were affixed to the skull and screws with dental cement. After surgery, rats were returned to their home cages and recovered for 1-2 weeks before beginning food restriction for the Set-Shift task.

### DBS-like electrical stimulation

Electrical stimulation(0.3mA, 130 Hz, 50 µs biphasic pulses) was delivered using a PC-controlled StimJim stimulator (Cermak et al., 2019). In prior work, locomotor effects were analyzed to ensure that any behavioral changes were not due to acute sensory or motor effects from the electrical current and that these changes were specific to stimulation at 130 Hz (Reimer et al., 2024). Rats received one hour of pre-task stimulation in an open field arena and throughout the duration of the task. For sham (“OFF”) stimulation sessions, rats were connected to cables without current delivery. Waveforms during stimulation sessions were monitored with an oscilloscope and a 1kOhm series resistor. To balance stimulation conditions across Set Shift sessions and to achieve adequate statistical power, each rat underwent 32 sessions total (8 per condition: OFF, bilateral, right, and left), in a pseudorandom, counterbalanced order. One rat developed seizure-like behavior after 28 sessions and two rats were only able to complete 31 sessions due to equipment malfunction. Because we sought to compare unilateral effects to the known bilateral RT decrease, 4 rats whose RTs increased with bilateral stimulation were excluded from the study. 3 of these 4 excluded rats had mistargeted electrodes, replicating prior findings (Reimer et al., 2024).

### Histology

Before implantation, electrodes were coated with fluorescent dye (DiI or DiO, Invitrogen, MA) for localization (DiCarlo et al., 1996). Rats were deeply anesthetized with isoflurane, euthanized (Euthasol, 100 mg/kg), and transcardially perfused with PBS followed by 4% paraformaldehyde. Brains were then extracted, post-fixed in PFA, cryoprotected in sucrose, and sectioned at 20 µm on a cryostat. Electrode placements were verified using images from a fluorescent microscope (Keyence) that were overlaid onto a rat brain atlas (Figure 2B). Placements were considered on-target if the electrode tips were within the mid-striatum boundaries defined by previous work (Reimer et al., 2024) (anterior-posterior range: +0.5-+2.5).

### Statistical Analysis

#### Design and Statistical Power

We used a power analysis (G*Power 3.1) to pre-determine sample size and number of sessions when analyzing RT, accuracy, and omissions data with generalized-linear models (GLMs). A GLM is functionally equivalent to a repeated-measures ANOVA, with within-factor effects. To achieve 80% power with an effect size of f = 0.25 and a Bonferroni-corrected alpha = 0.0167, we required 9 animals total. We therefore used a final sample of 10 rats (5 male, 5 female). Although we included sex as a regressor in some models, this study was not powered for sex differences, so these analyses should be considered exploratory.

#### Statistical Analysis of Set-Shift Behavior

We analyzed Set-Shift behavior in MATLAB (R2021b) using GLMs. GLMs are well-suited for repeated-measures data (different stimulation conditions in the same animal) and the non-Gaussian RT distribution, and are consistent with our prior work (Reimer et al., 2024). Our primary GLM included fixed effects for Stimulation, SessionNumber (1-32), and RuleType (1-side, or 0-light), with random effects for Chamber, Rat, and SessionID. The SessionID random effect was added to account for correlation among trials from the same session. Omission trials, where the rat did not respond during the 3 s trial window, were excluded from all RT and accuracy analyses. The general form of the GLM was therefore: *DependentVariable ∼ Stimulation+SessionNumber+RuleType+(1|Chamber)+(1|Rat) + (1|SessionID)* (Figures 1-2, 4).

We designed additional exploratory GLMs to investigate possible sex differences (Figure 3), that included interaction terms for Stimulation and Sex, Sex and SessionNumber, and Sex and RuleType, and the same random effects as above.

**Figure 3:**
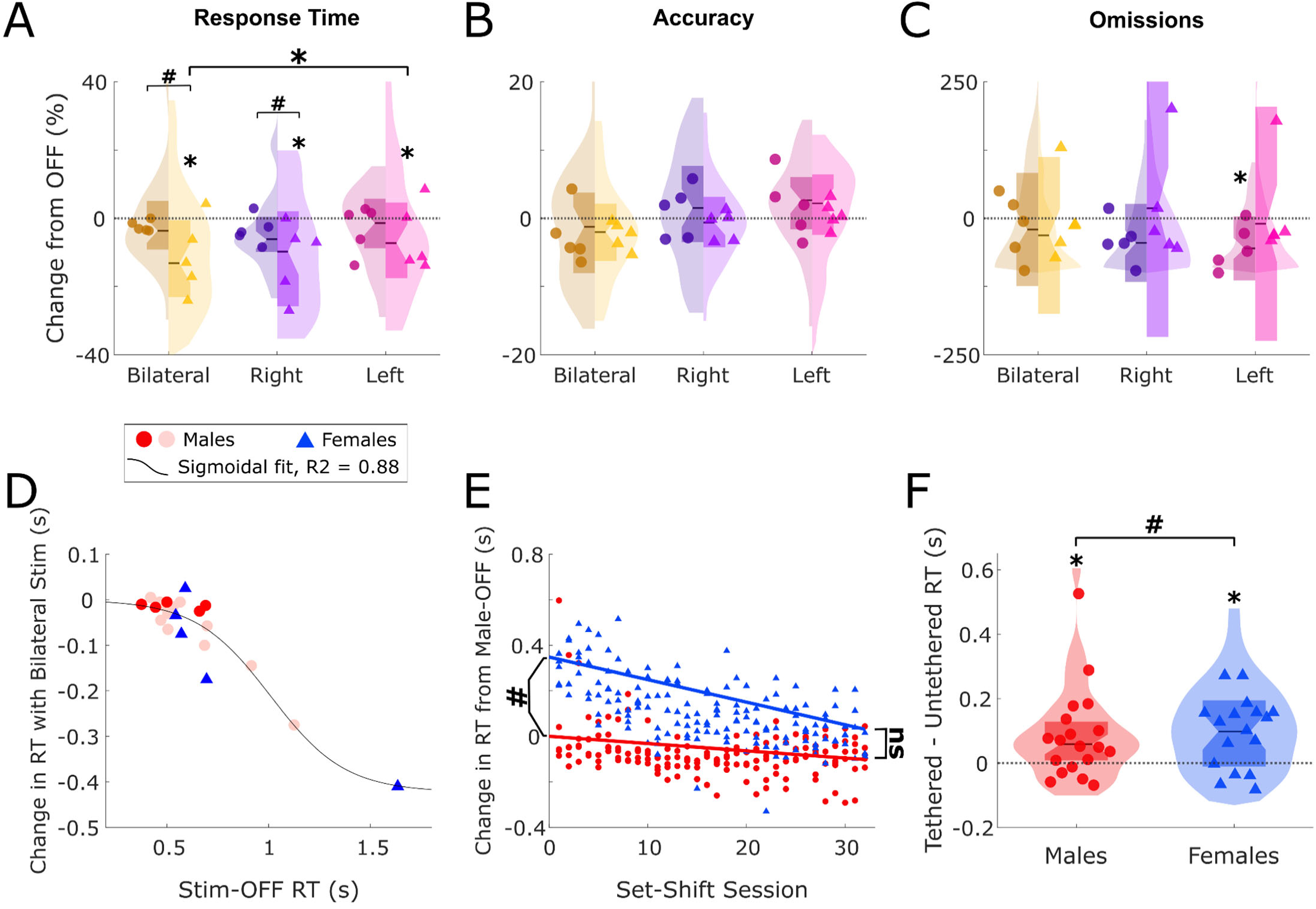
Sex differences in stimulation effects are explained by variables unrelated to cognitive control. A) Change in RT with bilateral (yellow) or unilateral (purple and pink) stimulation as percent-change from OFF RT, split by sex. Distributions quantify median RTs across Set-Shift sessions for males (left side of plots) and females (right side) separately, and individual scatter points represent the median RT of each rat (males = circles, females = triangles). **\*** indicates significant stimulation-condition intercept and # indicates significant stimulation-condition:sex intercept from gamma-distributed GLM B) Change in accuracy with stimulation as percent-change from OFF, split by sex. Distributions quantify the mean accuracy percentage across sessions for males and females separately, and scatter points represent the mean accuracy percentage of each rat with bilateral or unilateral stimulation. C) Change in trial omissions with stimulation as percent-change from OFF omissions, split by sex. Distributions quantify mean omission rates across Set-Shift sessions for males and females separately, and scatter points represent mean omission rates of each rat. D) Ceiling effect on the reduction in RT with bilateral stimulation. Plotted is each rat’s median reduction in RT with bilateral stimulation plotted against their OFF median. Males are indicated by red circles (light red indicates males from the mid-striatal cohort in (Reimer et al., 2024) and females are indicated by blue triangles. This data is fit by a sigmoid function (black line, R^2 = 0.88), where rats with faster OFF RTs had smaller reductions in RT with bilateral stimulation. E) Change in RT over the course of 32 Set-Shift Sessions by sex. Scattered is individual session mean RT, with the male-intercept and any random variation (male and female) removed. Plotted lines represent change in RT with session number relative to male OFF (intercept removed). F) Change in response time from cable tethering in males (red, left) and females (blue, right). A separate cohort of rats (N = 36) performed Set-Shift while untethered or tethered to a stimulation cable (without stimulation). Distributions quantify the median change in response time across sessions and scattered points represent each individual rat’s median change in response time (tethered - untethered).

P-values for GLM coefficients and combinations of terms (e.g. sex-related differences in rule-type accuracy) were computed using the *coefTest* MATLAB function. For our 3 main measures (RT, accuracy, omissions), we used a Bonferroni-corrected significance threshold of p < 0.0167. We then multiplied raw p-values by 3, and reported the corrected values (< 0.05, Figures 1, 2, 3A-C). Other secondary analyses used a standard p < 0.05 threshold (Figures 3E-F).

#### Reinforcement Learning-Drift Diffusion Model of Set-Shift Behavior

To provide more evidence that unilateral stimulation can replicate bilateral effects, we used a reinforcement learning-drift diffusion model (RLDDM) designed in prior work (Reimer et al., 2024). This model combines a drift-diffusion model that predicts RTs and choices based on latent decision-making variables, with a reinforcement learning model that captures choice variation based on reward history (Figure 4A). During each trial, the RLDDM assigns learned values to both nose pokes according to location and illumination. The difference in values between the left and right nose pokes is then leveraged to make a choice based on a drift-diffusion process. The difference in total value informs the drift rate (rate of evidence accumulation) and the difference in value between the left and right sides alone informs the bias (initial evidence favoring one side over the other). The total amount of evidence required to make a decision is the boundary separation. Once a choice is made, the values of the sides and light update based on the reinforcement learning process. After validating model fit and accuracy with posterior predictive checks, we determined the effect of stimulation on model parameters using the region of practical equivalence (ROPE). For a detailed explanation of these methods see (Reimer et al., 2024).

**Figure 4:**
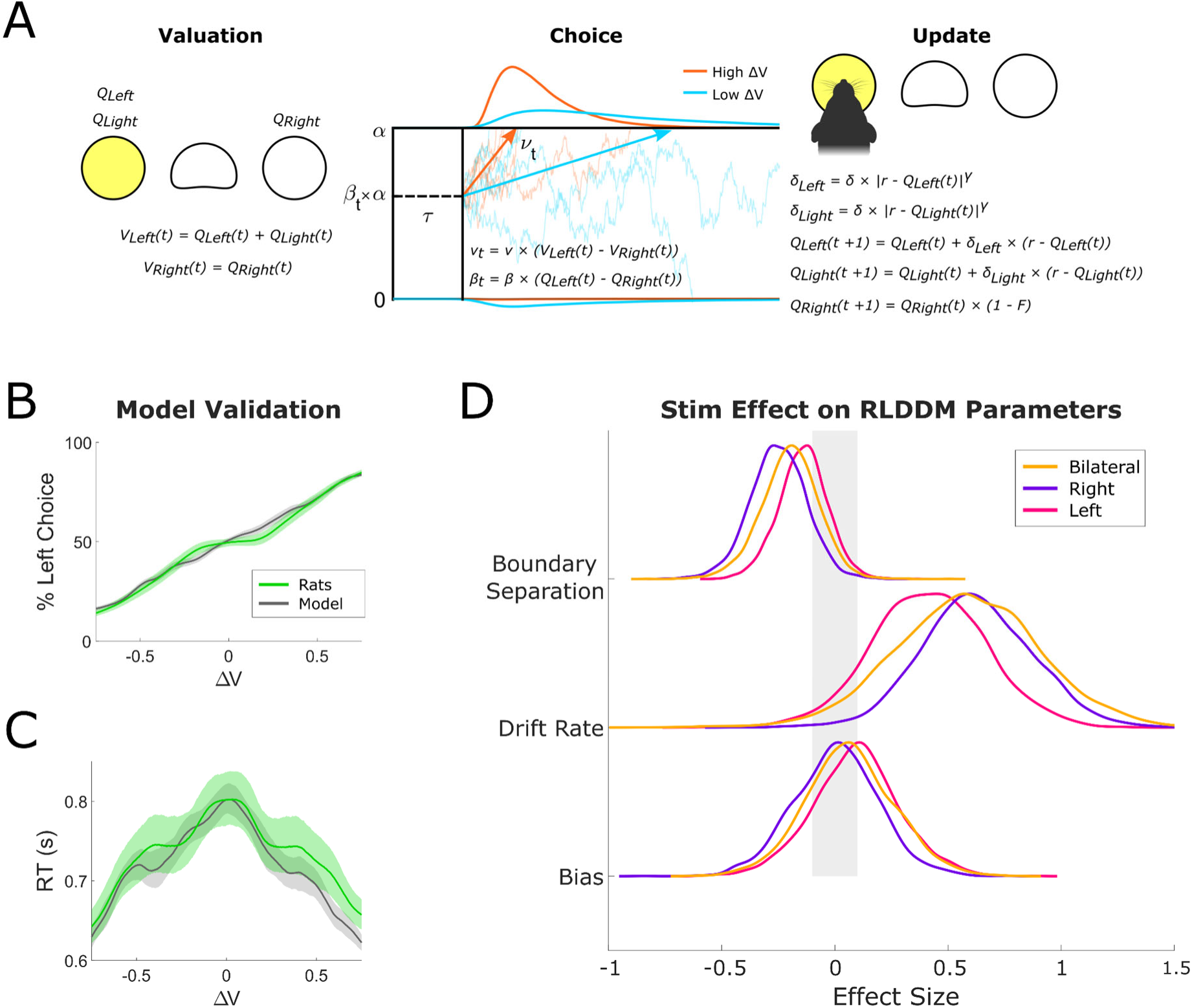
Stimulation Effects on Set-Shift Behavior Explained by RL-Drift-Diffusion Modeling. A) Schematic of the Reinforcement Learning-Drift Diffusion Model (RLDDM) to fit to Set-Shift behavior. The RLDDM first assigns values (V) to the ports based on its underlying estimates (Q) of the current value of the sides (left, right) and the light. The model then chooses a port based on a drift-diffusion process described by non-decision time τ, drift rate ν, and decision boundary α. The difference in total value informs the drift rate, while the difference between the sides alone informs the bias ß. Trials with high value differences are likely to have shorter RTs and more consistent choices (red vs. blue curves above). After the port choice, values are updated based on a reinforcement learning process, with a non-linear scaling factor γ to diminish small errors and accentuate large ones. The value of the unchosen option decays according to a forgetfulness factor F. The model was fit hierarchically to the data from all rats, allowing individual-specific estimates of model parameters and their change with stimulation. B) – C) Posterior predictive simulations of Set-Shift behaviors for models (gray) and actual rat behavior (green) over 4000 hierarchical posterior draws. Shading shows the 95% highest density interval across rats and trials. B) depicts side choice and C) depicts RTs. See Methods and (Reimer et al., 2024) for details. D) Distributions of the effect of stimulation on RLDDM model parameters. The grey shaded area represents the Region of Practical Equivalence (ROPE) for a null effect (effect size <0.1). Bilateral stimulation (gold line) affects boundary separation, drift rate, and bias, replicating results from (Reimer et al., 2024). These parameters were similarly altered by right (purple) and left (pink) unilateral stimulation, although the mean effect of right stimulation on boundary separation and drift rate was numerically greater than the effect of left stimulation.

## Results

Male and female rats were tested on the Set-Shift task, which engages cognitive control by requiring rats to shift between a light rule and a side (spatial) rule (Figure 1A) (Darrah et al., 2008; De Oliveira et al., 2021). We first evaluated RTs, accuracy, and omissions on non-stimulated (OFF) Set-Shift sessions. As in prior work, RTs were similar between side vs. light rule trials (Figure 1B). Rats were more accurate on light rule trials compared to side rule trials, a known feature of this task (Figure 1C,*, p < 0.00001). Although trial omissions were very infrequent, rats omitted more on side rule trials compared to light rule (Figure 1D, *, p = 0.000017), in line with the accuracy difference between rules.

### Unilateral stimulation replicates improved cognitive control from bilateral stimulation

We implanted electrodes in the mid-striatum where stimulation related improvements in RT were previously found (Reimer et al., 2024). A total of 10 rats were successfully implanted and completed 8 sets of four session blocks with bilateral, unilateral left and right, and OFF stimulation (Figure 2A-B). All three stimulation conditions significantly reduced RTs (Figure 2C, *, bilateral: p = 0.00009, right: p = 0.00024, left: p = 0.0193), with no significant differences between either unilateral condition and bilateral stimulation, although the improvement in RTs was numerically greatest from right stimulation. As in past work, no stimulation condition had any effect on accuracy relative to OFF (all *p*s > 0.05). However, accuracy was significantly higher for left compared to bilateral stimulation (Figure 2D, #, p = 0.0456). We additionally assessed stimulation effects on the probability of omissions. All three stimulation conditions reduced omissions, but this decrease was only significant for left stimulation (Figure 2E, p = 0.0309). Overall, this reduction in RTs and omissions, without decreased accuracy, signifies improved cognitive control from mid-striatal stimulation. The non-significant advantage for right-sided stimulation replicates a pattern in prior human data (Basu et al., 2023).

### Females and males have similar RTs and omissions, but different accuracy rates on SS

This was the first time females were tested on our version of Set-Shift. During Set-Shift training, we observed possible differences in Set-Shift behaviors and stim effects between the sexes, so we ran additional exploratory analyses (Figure 3). Females showed slower RTs than males for both rule types (light rule: p=0.0466, side rule: p = 0.0486) on OFF trials. Females were significantly less accurate than males but only in the side rule (p <0.00001). There were no sex differences in OFF trial omissions.

### Stimulation reduces RTs more in females than in males

Next, we assessed sex-related differences in stimulation effects (Figure 3). When considering each sex separately, the RT reduction from stimulation was driven primarily by females. In males, no stimulation condition significantly reduced RTs. Conversely, in females, all three conditions significantly reduced RTs compared to OFF (bilateral: p <0.00001, right: p <0.00001, left: p = 0.0099), and bilateral effects were greater than left effects (p = 0.0162) (Figure 3A). When comparing effects between sexes, bilateral and right stimulation reduced RTs to a greater degree in females compared to males (Figure 3A, #, bilateral: p = 0.0084, right: p = 0.0478), but left stimulation effects were similar between the sexes. There were no sex-related differences in stimulation effects on accuracy. For omissions, only males receiving left stimulation had a significant decrease (Figure 3C, p = 0.0291). The lack of a significant decrease in omissions in females is driven by one female who had increased omissions from all three stimulation conditions. There were no sex differences in stimulation effects on omissions.

The sex-related differences in stimulation effects on RTs do not necessarily mean that stimulation differentially alters cognitive control in females vs. males. Instead, it could be driven by other variables. For instance, because males were faster during OFF sessions, we hypothesized there may be a floor effect, limiting how much stimulation could reduce RTs. Confirming this, a plot of each rat’s OFF RT versus their RT change from bilateral stimulation showed a sigmoidal trend (Figure 3D, R^2^ = 0.88), where slower rats (predominantly females) showed larger improvements from stimulation.

Next, we plotted the change in RT over the 32 Set-Shift sessions for males and females, normalized to male OFF (Figure 3E). We found that females were significantly slower than males at session 1 (#, p = 0.0187), but became progressively faster over 32 sessions until this difference was eliminated (p > 0.05), while male RTs remained stable. This seems to point to sex differences in Set-Shift learning, but may also be driven by simple sexual dimorphism. We theorized that the physical effects of tethering to the stimulation cable might be greater in females, given that all males were at least 50g heavier than all females and generally have a larger body size. In a separate experiment (n = 36), we found that tethering slowed RTs of both sexes (*p*s < 0.00001), but to a greater degree in females than males (#, p = 0.0277) (Figure 3F). This suggests that the gradual improvement in female RTs observed during the stimulation experiment may reflect habituation to the physical burden of tethering.

### Unilateral and bilateral stimulation similarly enhance decision making, improving cognitive control

Using a RLDDM developed in prior work (Reimer et al., 2024), we tested whether the same parameter shifts from mid-striatal stimulation are present in the current dataset and if they differ for bilateral vs. unilateral conditions (Fig 4A). The model accurately fit Set-Shift behavior (Fig 4B-C). In agreement with the previous study, bilateral stimulation affected two RLDDM parameters: boundary separation and drift rate (Fig 4D, gold). Left and right stimulation had similar effects on drift rate, but right stimulation affected boundary separation slightly more than left stimulation (Fig 4D, pink and purple). This stimulation-induced reduction in boundary separation implies that less evidence is required to make a decision, while the concurrent increase in drift rate represents faster evidence accumulation. Together, these parameter shifts are thought to indicate more efficient resolution of value differences, a hallmark of cognitive control. Unlike the initial bilateral experiment, stimulation did not affect bias, the parameter containing information about the value of each nose-poke location. However, the previously observed small increase in bias contributed minimally to the overall cognitive control effect (Reimer et al., 2024).

## Discussion

We validated and refined a rodent model of VCVS DBS, adding to our understanding of how it enhances cognitive control. We demonstrate that unilateral DBS-like stimulation of the mid-striatum improves cognitive control in rats, mirroring the effects of bilateral stimulation. This stimulation-driven improvement in cognitive control is indicated by faster RTs on the Set-Shift task without decreases in accuracy and is further supported by changes in drift diffusion model parameters consistent with more efficient decision-making. Additionally, we found that unilateral stimulation, particularly right-sided, may be preferable over bilateral stimulation to improve cognitive control without unintended behavioral effects. This challenges the prevailing assumption that bilateral DBS is the optimal therapeutic approach in most patients. It also agrees with past suggestions that unilateral DBS is sufficient to reduce symptoms in some patients (Sturm et al., 2003) and that unilateral may be preferable over bilateral DBS to avoid aversive effects or boost specific cognitive functions (Widge et al., 2016; Lin et al., 2021; Del Bene et al., 2024). These key results reproduce our earlier findings (Reimer et al., 2024) and expand our mid-striatal DBS model to account for laterality effects observed in humans, where right-unilateral VCVS stimulation best improved cognitive control (Basu et al., 2023).

Our results offer insights into the neural components engaged by mid-striatal stimulation. The mid-striatal target contains axon projections from the medial PFC (mPFC) and local neurons, but we do not know which component stimulation engages to alter cognitive control. Human VCVS DBS is hypothesized to act primarily on axons within the ventral internal capsule, specifically those analogous to axons originating in rat mPFC (Heilbronner et al., 2016; Widge et al., 2019a). In both rats and humans, these axons are extensively interconnected across the right and left mPFC and bilaterally extend to the striatum and other basal ganglia structures (Gabbott et al., 2005). Therefore, unilateral mid-striatal stimulation may be able to improve cognitive control by acting on these axons, which in turn carry modulation to both mPFC hemispheres. Still, slight differences in the circuits engaged by bilateral, right, or left stimulation and their associated behavioral changes are probable given the known anatomical and functional lateralization in these regions (Gabbott et al., 2005; Lopez-Sosa et al., 2021; Korponay et al., 2022).

While we did not find any significant differences between the effects of different stimulation conditions, right stimulation resulted in the largest RT decrease. Additionally, bilateral stimulation worsened accuracy significantly more than left stimulation, but decreased omissions. Further, RLDDM analysis showed the greatest decrease in boundary separation from right stimulation, followed by bilateral and then left stimulation. Taken together, these results seem to indicate some lateralization in striatal functional circuitry and its response to DBS. For instance, detrimental crossing effects may be responsible for the decrease in accuracy and concurrent increase in omissions from bilateral, but not unilateral stimulation. This is supported by previous studies that found lateralized functional connectivity and cognitive effects from DBS delivered to other CSTC circuits (Waters et al., 2018; Del Bene et al., 2024). However, these differences also could be explained by other factors, like slight variation in electrode placement or improper stimulation parameters that could impair accuracy and worsen unintended motor symptoms. For example, our anterior-posterior range was ∼1mm, and more anterior electrode placement is closer to axonal CSTC components. Stimulation of more axonal vs. cell body components might be beneficial given the mechanistic hypotheses discussed above but may also generate off-target effects. More granular stimulation (e.g. with multiple electrode contacts) might be needed to detect more subtle laterality effects and differences in mid-striatal targeting.

In this study, we also explored sex differences in how mid-striatal DBS modulates cognitive control. The stimulation-induced improvement in RT was larger in females than in males. This lack of a significant stimulation effect in males was likely due to our study being underpowered (n = 5) to resolve the small effect size seen in previous work (n = 8) (Reimer et al., 2024). We found that this larger stimulation effect in females could be explained by two factors: the correlation between a rat’s baseline RT and their stimulation-induced RT improvement, and that smaller female rats are more physically impeded by cable tethering. Further testing with larger sample sizes and a more balanced task structure is needed to identify if true sex differences exist for DBS’s cognitive control effects. The effect of tethering and other variables like satiation and relative body size should be carefully considered in future experiments, particularly when dealing with small effect sizes or while assessing sex differences.

Beyond RTs and accuracy, we quantified the effect of mid-striatal stimulation on Set-Shift trial omissions, which occur when the rat does not respond within the three second trial window. Bilateral and both forms of unilateral stimulation reduced omissions. Recent work using DREADDs found that omissions during response inhibition are mediated through corticothalamic, but not corticostriatal connections (de Kloet et al., 2021). Those same corticothalamic projections, particularly those originating in mPFC, pass through our mid-striatal target and likely are engaged by stimulation.

Finally, we applied the RLDDM developed in a prior study to the current dataset to help explain how DBS alters behavior by modulating cognitive control. Replicating that prior work, bilateral stimulation increased drift rate (faster evidence accumulation) and decreased boundary separation (less evidence required to decide), and unilateral stimulation had similar effects. We were not powered to model males and females separately in this study, but in the future we hope to apply this RLDDM to assess how parameters change in females compared to males and determine which behavioral differences are linked to differences in cognitive control. Mice running a similar task show sex differences in decision-making during Set-Shift that may be amenable to RLDDM modeling (Glewwe et al., 2025). One limitation of this RLDDM version is that omissions are not modeled, which could skew parameter estimates (Leng et al., 2024). Overall, the similarity in parameter shifts from bilateral and unilateral stimulation reinforces the idea that unilateral DBS is sufficient to engage relevant CSTC cognitive control circuits.

This study enhanced our rodent model of human VCVS DBS by demonstrating that unilateral mid-striatal stimulation improves cognitive control as effectively as bilateral stimulation, potentially with fewer unintended behavioral effects. This aligns with findings from clinical studies. The cognitive control improvement was largest for right-sided stimulation, suggesting functional lateralization in CSTC circuits. We propose that unilateral stimulation works by acting on axons from both mPFC hemispheres, with potentially less detrimental cross-talk than bilateral stimulation. Future work with optogenetics, parametric optimization, and electrophysiological measurements in mPFC and striatum may help further detangle the variables that drive this cognitive control effect.

## Acknowledgements

This work was supported by MnDRIVE Brain Conditions and the US National Institute of Health (R01NS120851 to A.S.W.). E.M.S. was supported by the MnDRIVE Brain Conditions Neuromodulation Fellowship and by a T32 predoctoral fellowship (5T32DA007234). E.M.DvR. was supported by a National Science Foundation Graduate Research Fellowship (2237827). The opinions herein are fully those of the authors and do not represent the positions of any funding body.

## Author Contributions

E.M.S.: Conceptualization, Data curation, Formal analysis, Investigation, Methodology, Project administration, Supervision, Visualization, Writing-original draft

E. M. DvR.: Data curation, Methodology, Resources, Software, Validation, Writing-review & editing

J.P.B., M.C.B., M.E.M., & F.A.I.: Investigation, Validation, Writing-review & editing

A.E.R.: Conceptualization, Methodology, Resources, Validation

A.S.W.: Conceptualization, Funding acquisition, Supervision, Writing-review & editing

